# Hippocampal neural stem cells facilitate access from circulation via apical cytoplasmic processes

**DOI:** 10.1101/792192

**Authors:** Tamar Licht, Eli Keshet

## Abstract

Blood vessels (BVs) are considered an integral component of neural stem cells (NSCs) niches. NSCs in the dentate gyrus (DG) have enigmatic elaborated apical cellular processes that are associated with BVs. Whether this contact serves as a mechanism for delivering circulating molecules is not known. Here we uncovered a previously unrecognized communication route allowing exclusive direct access of blood-borne substances to hippocampal NSCs in defiance of an intact blood-brain barrier (BBB). BBB-impermeable fluorescent tracer injected transcardially is selectively uptaken by DG NSCs within a minute, via the vessel-associated apical processes. These processes, measured >30nm in diameter, establish direct membrane-to-membrane contact with endothelial cells in areas devoid of the endothelial basement membrane. Doxorubicin, a brain-impermeable chemotherapeutic agent, is also readily and selectively uptaken by NSCs and reduces their proliferation, which might explain its problematic anti-neurogenic or cognitive side-effect. The newly-discovered NSC-BV communication route explains how circulatory neurogenic mediators are ‘sensed’ by NSCs.

## Introduction

Neural stem cells (NSCs) serve as a source for new neurons in both the developing and adult brain. In the adult rodent brain, NSCs are confined into two locales, namely the subventricular zone (SVZ) of the lateral ventricles and the DG of the hippocampus. NSCs within the DG proliferate and differentiate into granule cells (GCs) eventually integrating into the existing network (van Praag et al., 2002). DG NSCs [also known as radial-glia-like cells (RGLs) (Bonaguidi et al., 2011)] are distinguished by a unique, tree-like morphology with soma embedded in the subgranular zone (SGZ), a major shaft extending through the adjacent granule cell layer (GCL) and terminating in a dense network of fine cytoplasmic processes spreading in the inner molecular layer (Gebara et al., 2016). The functional role of these terminal shafts (if any) has remained obscure.

Stem cell functioning in general, and of RGLs in particular, relies on microenvironmental support by various cellular and extracellular components, collectively referred to as the "stem cell niche". Blood vessels (BVs) are considered an integral, indispensable component of stem cell niches, including NSC niches (Licht and Keshet, 2015; Schildge et al., 2014). BV may impact stem cell performance by two different modes: via locally released blood-borne substances or via paracrinically acting factors elaborated by the endothelium (‘angiocrine factors’)(Rafii et al., 2016).

Adult hippocampal RGLs are mostly quiescent but undergo several rounds of cell division upon activation (Encinas et al., 2011; Pilz et al., 2018). Neurogenic rates are greatly affected by systemic cues, exemplified by exercise-enhanced neurogenesis shown to be mediated by systemic factors (Fabel et al., 2003; Ma et al., 2017; Moon et al., 2016). Likewise, heterochronic parabiosis experiments have demonstrated reciprocal influences on neurogenic rates mediated by systemic factors such as CCl11 or GDF11 (Katsimpardi et al., 2014; Villeda et al., 2011; Villeda and Wyss-Coray, 2013). Because hippocampal BVs (including those of the DG where RGLs reside) have a fully-functional BBB, the question arises how are circulating substances accessible to the brain parenchyma without a loss of BBB function?

Both SVZ-resident type B NSCs and DG-resident RGLs were shown to be intimately associated with BVs. In the SVZ, NSCs send a cellular process that wraps a capillary in the nearby striatum (Lacar et al., 2012; Licht and Keshet, 2015; Mirzadeh et al., 2008). In the DG, RGLs are physically associated with BVs at both the soma side (Filippov et al., 2003; Licht and Keshet, 2015; Palmer et al., 2000) and their apical side (Licht and Keshet, 2015; Licht et al., 2016; Moss et al., 2016). However, the functional significance of RGL-BV engagements is not well understood. Here we examined the proposition that these RGL processes might serve as a conduit through which blood-borne substances can be directly transferred to RGLs in face of a fully-functional BBB, thereby providing a mechanistic explanation of how can NSCs ‘sense’ circulating neurogenic mediators. Additionally, we show that systemically-injected doxorubicin (a BBB-impermeable DNA-intercalating agent used for non-cerebral tumors) can be uptaken by RGLs and provide a direct anti-mitotic mechanism for its anti-neurogenic effect.

## Results

### RGL processes contact endothelial cells in specialized basement membrane-free zones

The proposition that the thin processes extended by RGLs can serve as a bridge for transferring blood substances to RGL cell bodies requires that RGL-BV contact points are not separated by pericytes or by a basement membrane (BM) in which endothelial cells are usually invested and also constitutes an important component of the BBB (Thomsen et al., 2017). Previous studies that used Nestin-GFP to label RGLs (Filippov et al., 2003; Moss et al., 2016) were limited by the fact that mural pericytes are also labeled by this reporter (see Fig. 1a)(Nakazato et al., 2017), which makes perivascular pericytes and RGLs indistinguishable at the electron microscopy (EM) level. To overcome this shortcoming, we used Gli1-cre^ERT2^ mice [a mouse line in which inducible Cre is not expressed in DG cells other than RGLs (Ahn and Joyner, 2004)] crossed to Ai9 (TdTomato) reporter line. RGLs were then highlighted with anti-RFP antibody and DAB staining (Fig. 1b) and the tissue processed for EM imaging. Soma of DAB-labeled RGLs in the SGZ were readily discernible by their DAB dark granules (Fig. 1c). At the inner molecular layer in the vicinity of that soma, we identified capillaries that are tightly wrapped by extra-fine (up to 30nm in diameter) RGL processes labeled with DAB (Fig. 1 d-g).

**Fig. 1.**
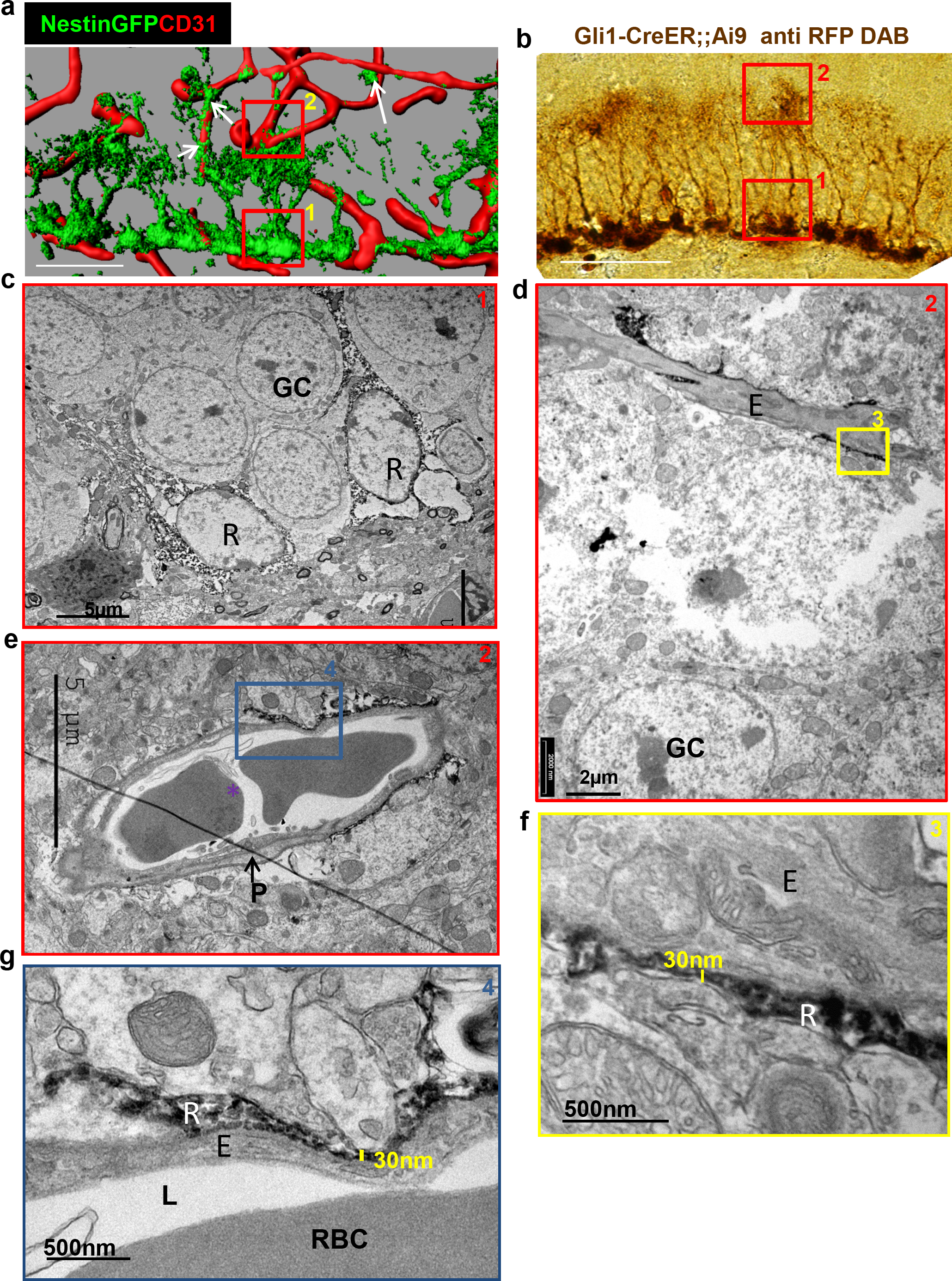
Fine apical processes of RGLs enfold capillary BVs in the inner molecular layer. **a** 3D reconstruction of the DG of Nestin-GFP mouse co-stained with CD31 for blood vessels. In this line, both RGLs and pericytes (arrows) are GFP-labeled. Please note the contact points of RGLs and blood vessels at both the basal side at the SGZ (1) and the apical side at the inner molecular layer (2). Scale bar, 50μm. **b** Gli1-Cre^ERT2^ crossed to Ai9 line (inducible TdTomato reporter) received tamoxifen three days before brain retrieval. Immunohistochemistry was done using anti-RFP antibody and DAB labeling. Note that only RGLs are labeled in the DG of this mouse line. (1) basal side. (2) apical side. Scale bar, 50μm. **c** Transmission electron microscopy (TEM) images taken at the SGZ (1) showing the soma of two DAB-labeled RGLs (R). The cytoplasm of those cells is detected by DAB dark granules. GC-granule cells. **d** and **e** TEM images of the inner molecular layer (2) showing two representative examples of a capillary wrapped with DAB-labeled RGL processes. **f** and **g** Higher magnification of (**d**) and (**e**) showing that RGLs processes can reach up to 30nm in thickness. R-RGL; GC-granule cell; E-endothelial cell, RBC-red blood cell; L-blood vessel lumen; P-pericyte.

The BM is visualized in EM images as a grayish 50-100nm-thick layer all around the abluminal aspect of endothelial cells (Fig. 2a). This component was found to thoroughly encircle DG capillaries of the hilus and outer molecular layer. Strikingly, however, zooming-in to endothelial-RGL processes contact points in the inner molecular layer revealed that engagement with BVs takes place in endothelial surfaces distinguished by the absence of a classical BM appearance, thus allowing for a direct membrane-to-membrane contact without the barrier imposed by the BM (Fig. 2b). Immunofluorescence for the endothelial BM component laminin also revealed a reduction in its expression at the RGL-BV contact point (Supplementary Fig. 1). The BV-RGL interface was also characterized by robust vesicular activity as many cytoplasmic and abluminal membrane-bound vesicles were detected (Fig. 3). In view of these findings, this specialized neurovascular unit (NVU) was tested for allowing the infiltration of systemically-derived molecules.

**Fig. 2.**
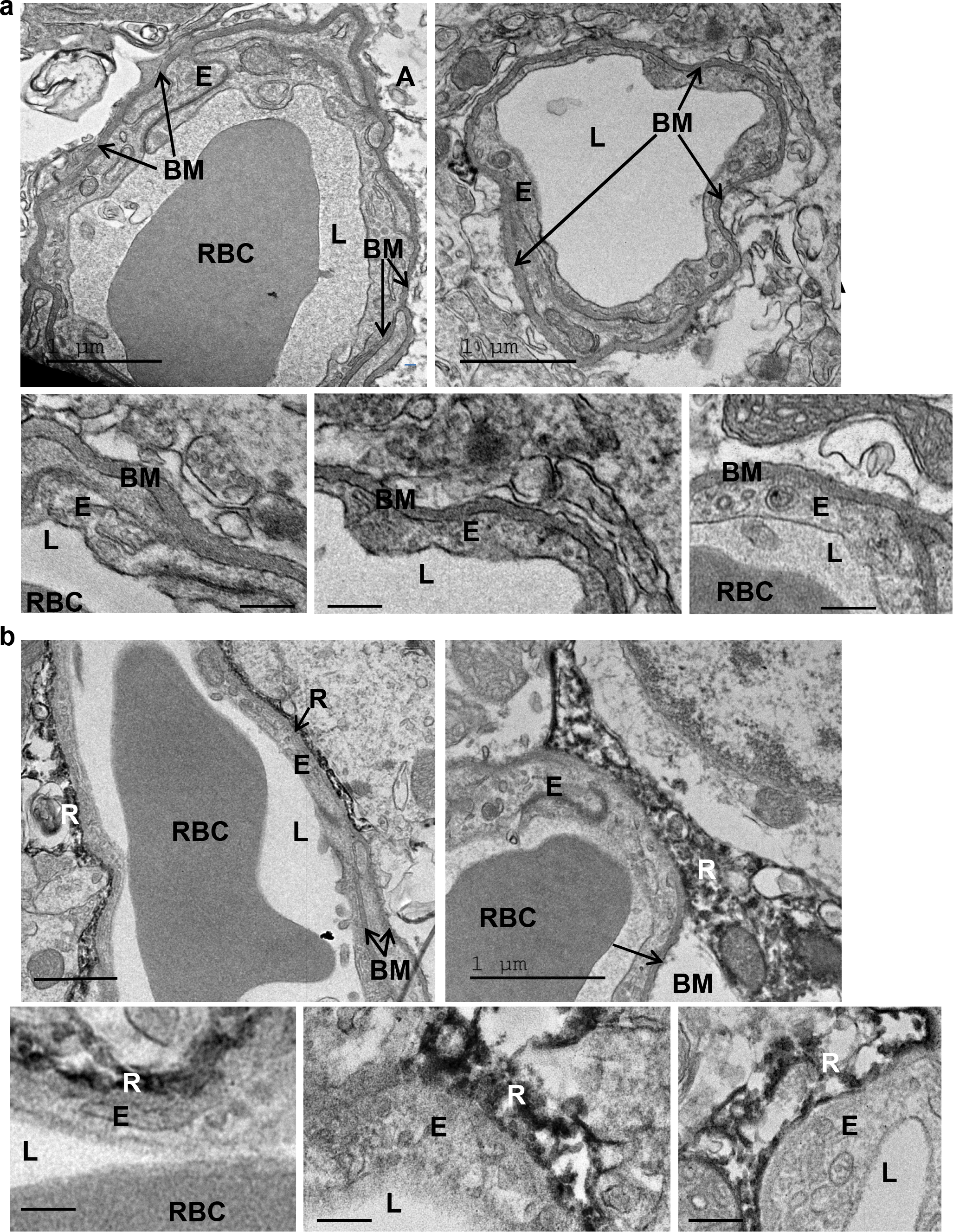
Lack of distinctive basement membrane at the contact points between RGLs and endothelial cells in the inner molecular layer. TEM images of the same samples as in Fig. 1. **a** Representative images of capillaries from the hilus area. Note BM covering the entire capillary abluminal membrane. Bottom– higher magnifications. Scale bars: top - 1μm, bottom-100nm. **b** Representative images of capillaries taken from the inner molecular layer that associate with DAB-labeled RGL. Note no characteristic BM at the contact point of RGL (R) with the endothelial cell (E). Scale bars: top - 1μm, bottom-100nm. R-RGL; E-endothelial cell, RBC-red blood cell; L-blood vessel lumen; BM-basement membrane.

**Fig. 3.**
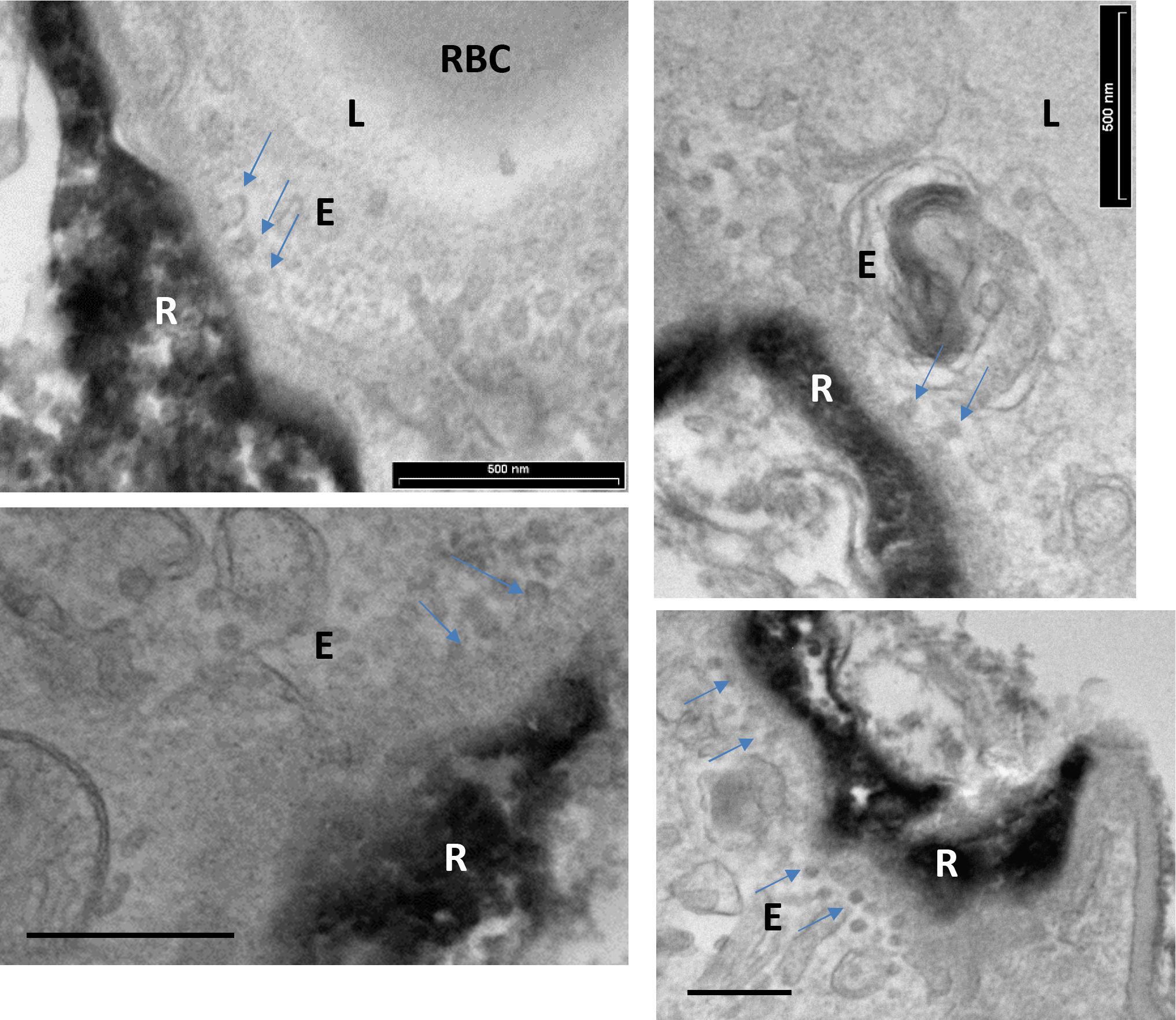
Vesicular activity at the BV-RGL interface. TEM images were taken from the same samples as in Fig. 1 and zoomed to BV-RGL contacts at the inner molecular layer. Arrows highlight abluminal membrane-connected and cytoplasmic endothelial vesicles. Scale bar, 500nm. E-endothelial cell, R-RGL, L-BV lumen, RBC-red blood cell.

### BBB-impermeable circulatory molecular tracer is uptaken exclusively by hippocampal RGLs

Circumventricular organs are known to have permeable vessels to allow for direct exposure of neurons to circulating components. To examine the possibility that other cells in the intact adult brain are ‘privileged’ in the sense of having access to blood-borne substances, we injected into the systemic circulation a fluorescently-tagged 10Kd dextran tracer previously shown to be fully contained in the lumen of blood vessels with a functional BBB (Ben-Zvi et al., 2014; Licht et al., 2015). The tracer (6mg/kg) was injected to the left ventricle of the heart (the tracer is rapidly cleared in the urine when injected to the tail vein) and the brain was retrieved one minute later and immediately fixed. Inspection of cryosections, co-stained with CD31 to highlight BVs, confirmed that - with the exception of the hypothalamus (a circumventricular organ) and choroid plexus vessels which are known to lack a functional BBB - the tracer is indeed fully retained in nearly all other cerebral BVs (Fig. 4a). The only cells accumulating the systemically-injected tracer were a small population of DG cells (Fig. 4a, top left). Remarkably, within the DG, tracer uptake is confined to a group of cells in which their soma is located in the SGZ, i.e., coinciding with the region where RGLs soma reside, and a major shaft is extended towards the molecular layer (Fig, 4b). In fact, both the location and unique morphology of tracer-labeled cells strongly suggested that these are DG RGLs.

**Fig. 4.**
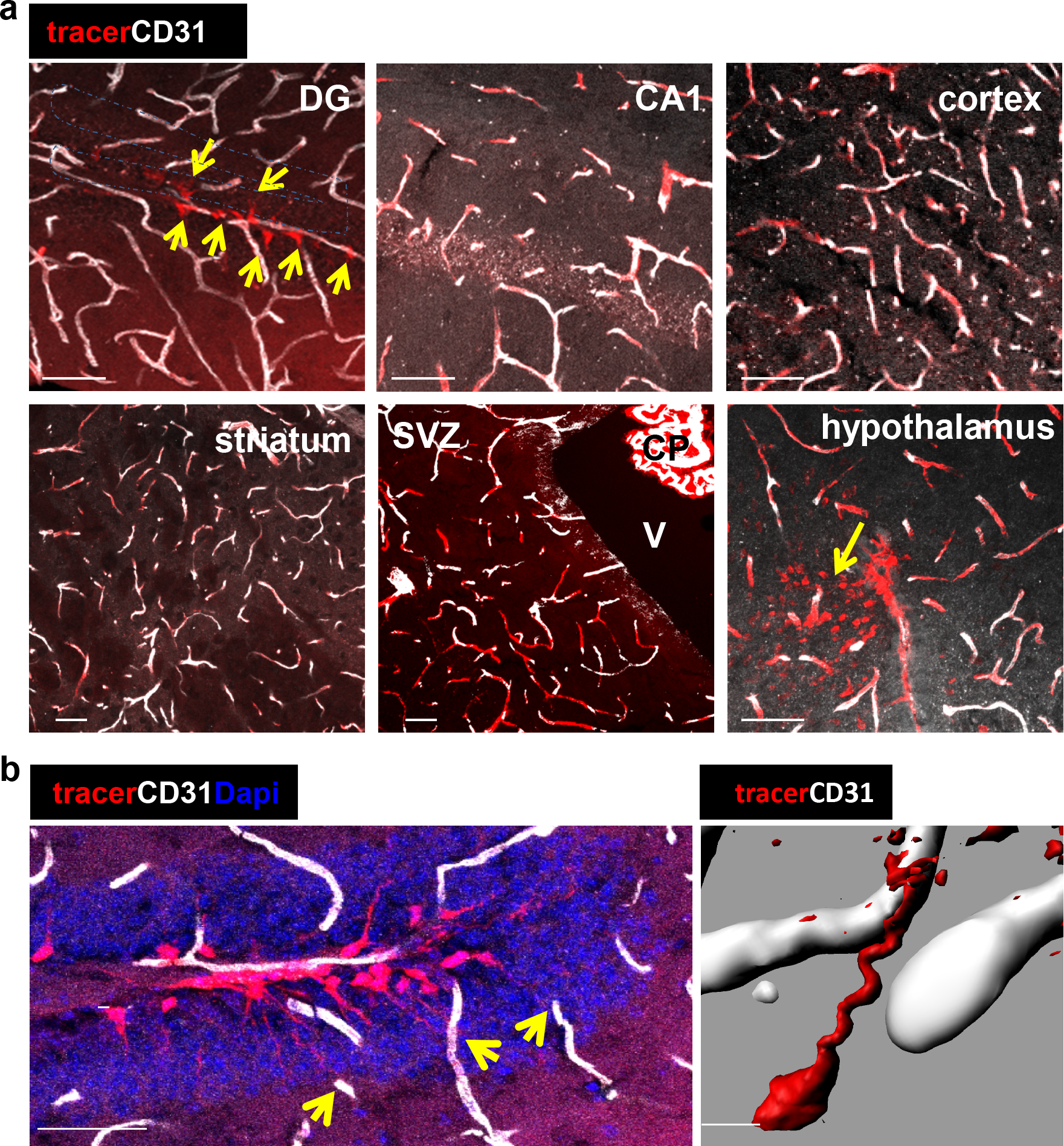
A unique population of cells in the SGZ is specifically labeled by systemically-injected 10Kd dextran tracer. **a** 10Kd dextran TRITC-labeled tracer was injected to the left heart ventricle of an anesthetized mouse. One minute later the brain was retrieved and immediately fixed. Sections (co-stained with CD31) of the indicated brain areas are presented. Tracer did not infiltrate into cells outside of the vasculature except for a specific population of cells in the SGZ of the DG (arrows), the CP and the hypothalamus (arrow). Scale bar, 50μm. V-cerebral ventricle. CP – choroid plexus. (**b** Representative high magnification image of the DG area in TRITC-10Kd dextran-injected animal co-immunostained with CD31. Labeled SGZ cells have a radial process that is associated with local blood vessels in the inner molecular layer (arrows). Scale bar, 50μm. Right: – 3d reconstruction of a dextran-labeled cell and a capillary illustrating mutual contact with no labeling of the extracellular space. Scale bar, 5μm.

We wished to determine whether these cells serve as a conduit allowing direct transfer from the circulation without extracellular leakage. We followed the trajectory of a tagged cell from a vessel-associated cellular process to soma. Considering its tortuous path, we used Imaris-aided 3D-reconstructions of serial confocal images. As shown in Fig. 4b (right), there is a clear continuum in tracer presence from the vessel through the major shaft to the soma with no sign whatsoever of extracellular spillage.

To verify the identity of the tracer-accumulating cells, TRITC-labeled 10Kd tracer was injected into the circulation of mice harboring a Nestin-GFP reporter transgene highlighting RGLs (here, in contrast with EM studies, they can be distinguished from pericytes by their tree-like morphology). As shown in Fig. 5a, due to their fineness, the tracer was barely visible in the tree-like processes and most of it accumulated in the major shaft and soma. We wished to re-confirm that RGLs thin processes also possess the tracer. For this purpose, we injected Nestin-GFP mice with biotin-labeled 10Kd dextran and visualized tracer distribution with Cy3-labeled streptavidin (this procedure was preferred to reduce interference of TRITC signal with GFP fluorescence). Colocalization of biotinylated tracer with nestin-GFP RGLs was evident in both the soma and the apical processes (Fig. 5b).

**Fig. 5.**
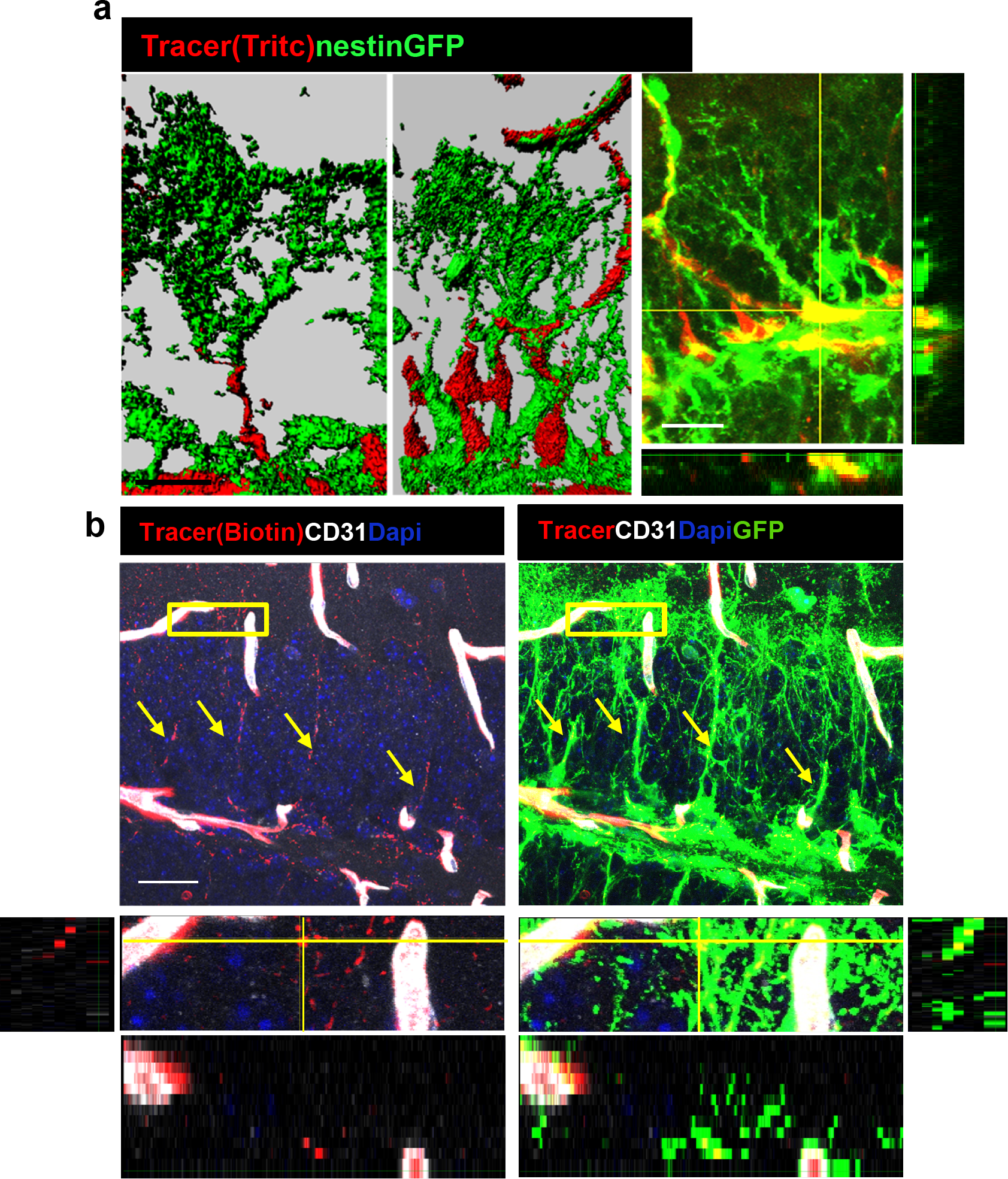
DG RGLs uptake a 10Kd systemically-injected dextran tracer. **a** TRITC-labeled 10Kd tracer was injected into a Nestin-GFP mouse. 3D reconstruction (left) and confocal z-plane (right) images showing accumulation of the tracer in the soma and major shaft of RGLs. Scale bar, 20μM. **b** Biotin-labeled 10Kd tracer was injected as in (**a**). Sections were stained for CD31 and tracer was highlighted by cy3-labeled streptavidin. Note tracer distribution throughout all RGL processes and soma (arrows), although less strong than the fluorescent tracer. The inset demonstrates the localization of tracer within GFP+ RGL fine apical processes. Scale bar, 20μM.

### Direct uptake of the chemotherapeutic agent doxorubicin by RGLs explains its anti-neurogenic side effects

Doxorubicin (dox) is a widely-used chemotherapeutic agent of the Anthracyclines family of DNA-intercalating agents. While inaccessibility of this small molecule (543Da) to the brain parenchyma (Ohnishi et al., 1995; Treat et al., 2007) has precluded its use for treating brain tumors, its use against other tumors is often associated with significant cognitive impairment, also known as “chemo brain” (reviewed by (El-Agamy et al., 2019)). Findings in rodents showing that dox have a dramatic negative effect on DG neurogenesis (Christie et al., 2012; Janelsins et al., 2010; Kitamura et al., 2015; Park et al., 2018)) prompted us to examine whether this seemingly paradoxical situation could be explained by direct selective uptake of systemically-administered dox by RGLs. To this end, an experimental protocol similar to the one used for following tracer trafficking was used, except that dox (6mg/kg) replaced dextran injection. Here we took advantage of the fact that dox have the same fluorescent properties as commonly used red fluorophores thereby enabling its direct visualization in cell nuclei in PFA-fixed brain sections. The fact that systemically-injected dox does not cross to brain parenchyma was validated by showing that in all brain regions known to have a functional BBB, dox was detected only in endothelial cell nuclei (exemplified for the cortex and CA1 region of the hippocampus in Fig. 6a), whereas in areas in which the BBB is inactive, dox-labeled nuclei were also readily detected outside the vasculature (shown for the choroid plexus and the median eminence of the hypothalamus in Fig. 6a). An outstanding exception was dox uptake by RGLs, evidenced by significant labeling of RGL nuclei within 1-2 minutes from its injection into the circulation of nestin-GFP mice where co-labeling with GFP confirmed the RGL identity of dox-uptaking cells (Fig. 6b and c). We next examined whether dox (being an anti-mitotic agent) has a negative effect on RGL proliferation. For this purpose, we injected dox at the dose of 50mg/kg into the peritoneal cavity of Nestin-GFP mice (three injections, every other day) followed by three injections of BrdU (50mg/kg, every 12 hours) to tag proliferating cells (Fig. 7a). Dox treatment did not have an effect on the total numbers of Nestin+ RGL population (Fig. 7b, right), as the majority of those cells are not proliferating. We immunostained for both the proliferation marker Ki67 (Fig. 7b) and for BrdU (Fig. 7c) to detect the proliferating RGL populations. In both cases, the numbers of proliferating RGLs have decreased significantly following dox treatment (Fig. 7b and c). We, therefore, suggest a direct antimitotic effect of dox as a presumable explanation for reduced neurogenesis reported in rodents treated with dox and as a possible factor in the “chemo-brain” side effect of cancer patients.

**Fig. 6.**
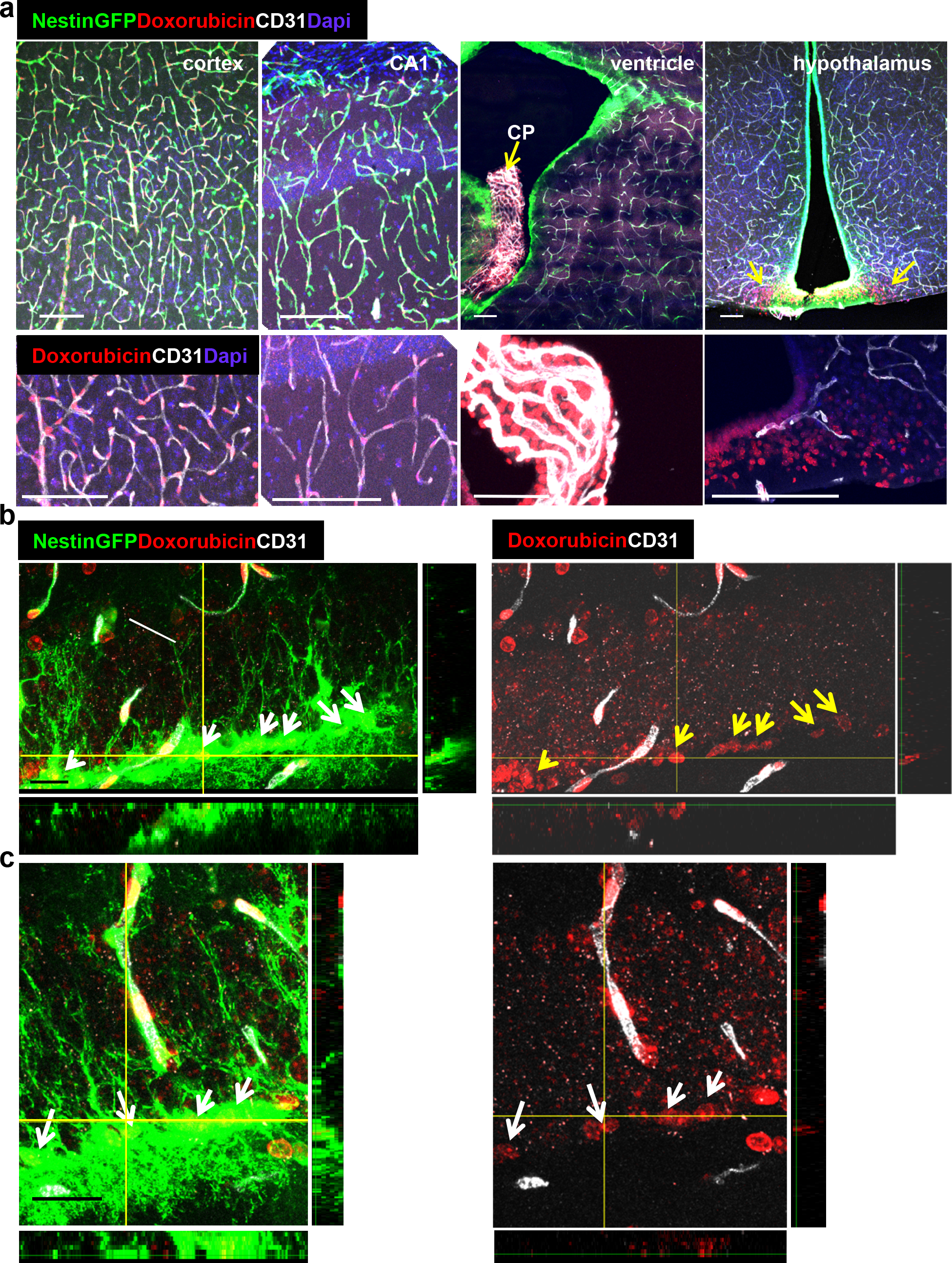
Doxorubicin has limited permeability through BBB capillaries but is being uptaken by RGLs. Doxorubicin (200μg) was injected into the left heart ventricle of Nestin-GFP mice. The brain was retrieved two minutes later. Doxorubicin is integrated into the DNA and detected by self red spectrum fluorescence. **a** Immunofluorescence images co-stained with CD31 showing integration of doxorubicin to the nuclei of various brain areas. In areas with intact BBB such as the cortex and the CA1, doxorubicin integrates solely to capillary nuclei. In areas with non-barriered endothelium such as the choroid plexus and the hypothalamus, doxorubicin is detected externally to blood vessels (arrows). Scale bar, 100μM. **b** and **c** Two representative high-magnification images of the DG demonstrating the integration of doxorubicin to the nuclei of GFP+ RGLs. Z-plane projections are shown. Scale bar, 20μM.

**Fig. 7.**
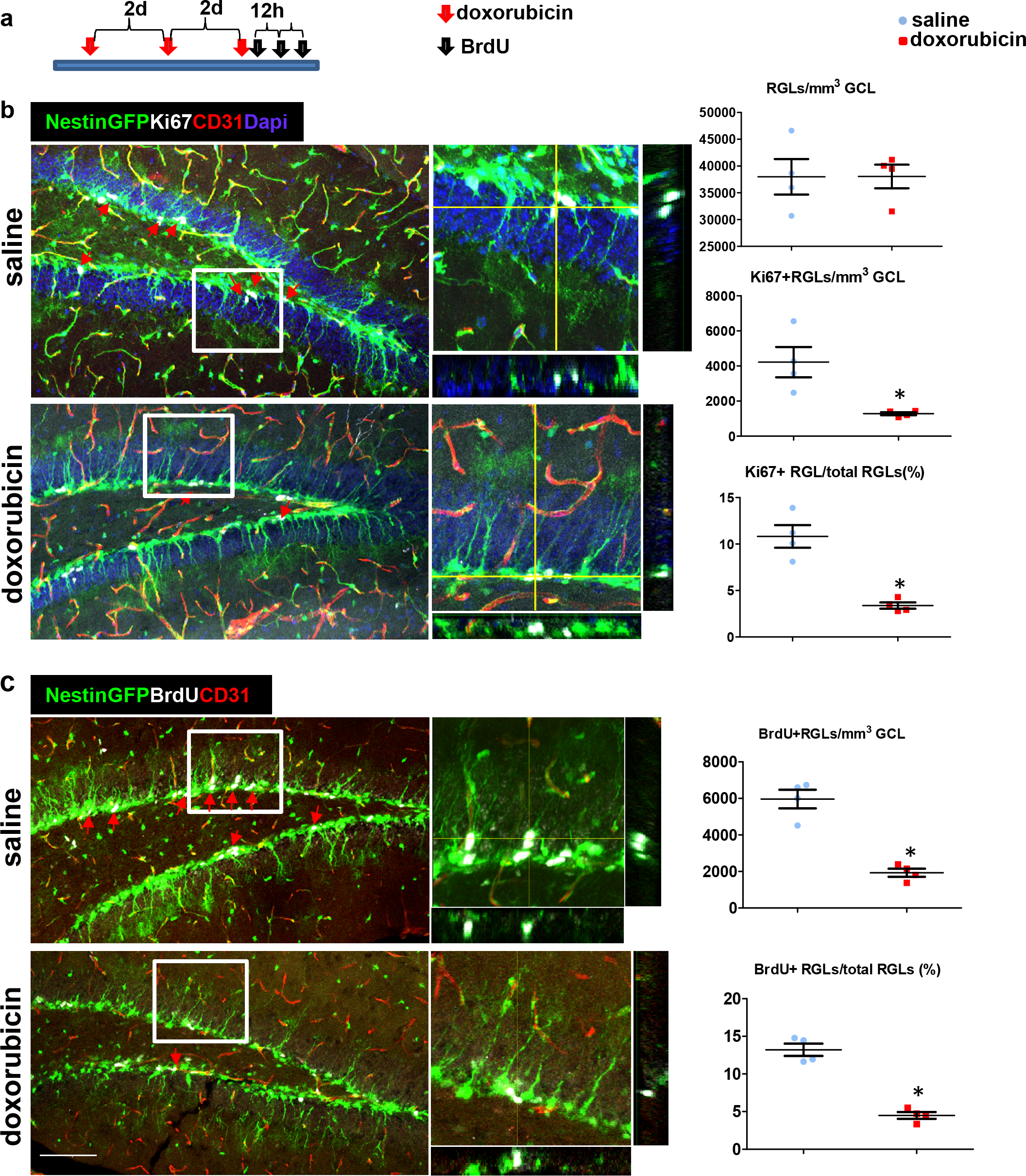
Doxorubicin treatment reduces the proliferation rates of Nestin-GFP^+^ RGLs. **A** Experimental protocol. Doxorubicin (5mg/kg) or saline were intraperitoneally (I.P) injected to Nestin-GFP mice every other day (3 injections). BrdU (50mg/kg) was injected I.P. to all animals 3 times at 12h intervals before brain retrieval. **b** Immunofluorescence images of nestin-GFP DG co-stained with Ki67 (a proliferation marker) and CD31. Arrows highlight Ki67+GFP+ cells that were verified to have a radial cytoplasmic process (see enlarged z-projection insets). Right: quantification of the total Nestin-GFP RGL population density (top, t(6)=−0.016 p=0.988), Ki67+ RGLs density (t(6)=3.398 p=0.015) and the percentage of Ki67+ RGLs among total RGLs (t(6)=5.923 p=0.001). **c** DG sections were co-stained with BrdU and CD31. Arrows highlight BrdU+GFP+ cells that were verified to have a radial cytoplasmic process (see enlarged insets). Right: quantification of BrdU+ RGLs density (t(6)=7.285 p=3.4*10^−4^) and the percentage of BrdU+ RGLs among total RGLs (t(6)=9.331 p=8.58*10^−5^). Scale bar for all images, 100μM.

## Discussion

Findings reported here showing that RGL-type NSCs in the adult hippocampus have a ‘privileged’ direct access to circulating blood molecules provide a possible mechanistic explanation to the enigmatic situation of NSCs ‘sensing’ the presence of particular blood components and responding by adjusting neurogenic rate despite the existence of a fully functional BBB. Circulatory molecules were shown to increase (or decrease) neurogenesis include specific cytokines whose circulatory levels change with age, as well as blood components accumulating in plasma following physical exercise and shown to upregulate DG neurogenesis (Castellano et al., 2017; Cooper et al., 2018; Fabel et al., 2003; Leiter et al., 2019; Ozek et al., 2018; Smith et al., 2015; Villeda et al., 2014). It is likely that additional physiologic cues impacting DG neurogenesis are also mediated by systemic factors making use of this direct gateway to DG RGLs.

Our finding showing that the transfer of blood-borne molecules to RGL cell bodies takes place through RGL’s thin cytoplasmic processes ascribes a first known function to the obscure terminal arbor of RGLs. Noteworthy, unlike RGLs, SVZ NSCs failed to uptake injected molecules (Fig. 4a) despite the fact that they also contact BV via a single extended process(Mirzadeh et al., 2008). Previously, we and others have shown that cytoplasmic processes extended by RGLs make direct contacts with nearby BVs (Licht and Keshet, 2015; Licht et al., 2016; Moss et al., 2016). Moreover, we showed that VEGF-promoted BV rejuvenation leads to remodeling of the terminal RGL arbor and engagement with even more remote capillaries (Licht et al., 2016). Yet, the functional significance of these physical RGL-BV associations has remained unknown until now.

We show, using EM images, that the fine processes of RGLs in the DG make direct contact with the membrane of nearby endothelial cell in the inner molecular layer. This contact takes place in areas with modified/reduced endothelial BM. We also highlighted extensive endothelial vesicular activity at the engagement points. These findings suggest that systemic molecules are delivered via vesicular transcytosis directly to the membrane of RGLs. Further studies are required to verify how the transfer of molecules takes place and whether this process is discriminative with regards to specific molecules.

Indiscriminative uptake may explain apparent RGL vulnerabilities to BBB-impermeable molecules. A notable, clinically-relevant example is that of the BBB-impermeable chemotherapeutic agent doxorubicin. A major side effect of dox is a debilitating cognitive decline affecting a considerable fraction of treated cancer patients (El-Agamy et al., 2019) and animal studies have shown that treatment with dox is manifested in impeded neurogenesis (Christie et al., 2012; Janelsins et al., 2010; Kitamura et al., 2015; Rendeiro et al., 2016). However, it has been difficult to reconcile the adverse cerebral side effects of dox with the fact that it is a substrate for p-glycoprotein pumps and is not available to the brain. Our finding that systemically-injected dox is readily and efficiently uptaken by DG RGLs provides a more compelling explanation to this seeming paradox than previous suggestions arguing an indirect mechanism (Janelsins et al., 2010; Kitamura et al., 2015). We show here that the treatment by dox decreases proliferating RGL numbers thus argue for a direct anti-mitotic mechanism.

We conclude that this unique NVU pathway may serve as a gateway for molecules that need to bypass the barrier or a "hole in the fence" allowing for penetration of molecules that should not enter the brain. We believe that the new mode of delivery of systemic molecules to RGLs uncovered by this study, not only provides a mechanistic explanation to systemic influences on adult hippocampal neurogenesis but may also provide new means of interventional modulations of this important process.

## Methods

### Mice

All animal procedures were approved by the animal care and use committee of the Hebrew University. Transgenic mouse lines that were used in this study (all of C57/Bl6 background): Ai9 and Gli1-cre^ERT2^ lines were purchased from the Jackson Laboratories (strains 007909, 007913). Nestin-GFP line was obtained from Prof. Grigori Enikolopov, CSHL (Mignone et al., 2004). Both males and females aged 2-3 months were used. Animals were grown in SPF housing conditions (4-6 animals per cage) with irradiated rodent food and water ad libitum and 12h light/dark cycle. Tamoxifen (Sigma T5648, 40mg/ml in sunflower seed oil) was administered orally once daily for 2-3 days at a dose of ~8mg/animal. BrdU (Sigma B5002) was dissolved in saline and injected I.P at 50mg\kg every 12h (3 injections).

### Tracer and doxorubicin injection

The following dextran tracers were used (all are lysine fixable): Tritc-labeled 10kD dextran (Molecular Probes, D1817), biotin-labeled 10Kd dextran (D1956). Doxorubicin was obtained from Sigma-Aldrich (44583). All were dissolved in saline to 2mg/ml and 100μl (6mg/kg) was injected into the left heart ventricle 1-2 min before brain retrieval. For chronic use, doxorubicin was injected I.P. at 5mg/kg every other day, three times.

### Tissue processing for immunofluorescence

Brains were fixed by perfusion and immersion in 4% PFA for 5 hr, incubated in 30% sucrose, embedded in OCT Tissue-Tec and cryosectioned to 50μm floating sections. Coronal slices from all aspects of the rostral-caudal axis were examined. For BrdU staining, brain slices were incubated for 2h in 50% formamide/2xSSC at 65°C, 30min in 2N HCl at 37°C and 10min in 0.1M boric acid pH8.5. Staining was done as described (Licht et al., 2010) with the following: anti BrdU (Serotec 1:400 PRID: AB_323427), anti-CD31 (BD 1:50 PRID: AB_393571), anti Laminin (Thermo Scientific 1:200, RRID:AB_60396), Anti Ki67 (Thermo Scientific 1:200, PRID:AB_10979488), anti-GFP (Thermo Scientific 1:200, PRID: AB_221570) and anti-RFP (Abcam 1:400, PRID AB_945213). Cy5 and Cy3 anti-rat, as well as Cy3 streptavidin, were obtained from Jackson Immunoresearch (dilution 1:400). Sections were mounted with Permafluor mounting medium (Thermo Scientific, TA-030-FM) with Dapi (Sigma, D9542). Confocal microscopy was done using Olympus FV-1000 on 20X and X60 lenses and 1.46μm distance between confocal z-slices. 3d reconstruction images were processed by Bitplane IMARIS 7.6.3 software using the Isosurface function.

### Cell proliferation quantification

6-7 confocal z-stack images per animal were quantified (n=4). The volume of the GCL was calculated by measuring the GCL area in the image (using Olympus FV-10 viewer) and multiplying the measured area by the number of z-axis slices and the distance between slices (1.46μm). Cells within the GCL (Ki67/CldU/GFP) were counted manually by a blind experimenter (using Olympus FV-10 viewer). Z-plane projections used to verify double labeling. Student’s T-test (SPSS) was used to compare between groups, assuming a normal distribution and unequal variance.

### Tissue processing for electron microscopy

Brains of Gli1-Cre^ERT2^;;Ai9 mice (3 days post tamoxifen induction) were transcardially perfused with PBS solution (5mL over 1min) and then fixed (50mL over 10min, 4% paraformaldehyde (wt/vol); 0.1% glutaraldehyde (vol/vol) in 0.1M phosphate buffer; and kept at 4 °C for 24 h. The brain was then washed in PBS solution, and cut in 50-μm coronal sections with a vibratome (Leica). For the immunoperoxidase process, the tissue was washed 10min in 3% H2O2 solution, blocked by two washes in 0.5% BSA and the sections were incubated overnight in the primary antibody (anti RFP, Abcam 1:400, PRID AB_945213) in 0.1% BSA-PB with shaking at 25 °C. Next, sections were washed in PBS solution and incubated in the secondary antibody (anti-rabbit HRP, Vector MP-7451-50) shaking for 2 h at 25 °C. Sections were then washed in PBS solution and incubated in DAB/plus chromogen solution (Abcam ab103723) for 6-10min, followed by subsequent washes in PBS solution (30min). Stained brain slices were transferred back to the fixative solution, were rinsed 4 times, 10 minutes each, in cacodylate buffer and post-fixed and stained with 1% osmium tetroxide, 1.5% potassium ferricyanide in 0.1M cacodylate buffer for 1 hour. Tissue was then washed 4 times in cacodylate buffer followed by dehydration in increasing concentrations of ethanol consisting of 30%, 50%, 70%, 80%, 90%, 95%, for 10 minutes each step followed by 100% anhydrous ethanol 3 times, 20 minutes each, and propylene oxide 2 times, 10 minutes each. Following dehydration, the tissue was infiltrated with increasing concentrations of Agar 100 resin in propylene oxide, consisting of 25, 50, 75, and 100% resin for 16 hours each step. The tissue was then embedded in fresh resin and let polymerize in an oven at 60°C for 48 hours. Embedded tissues in blocks were sectioned with a diamond knife on a Leica Reichert Ultracut S microtome and ultrathin sections (80nm) were collected onto 200 Mesh, thin bar copper grids. The sections on grids were sequentially stained with Uranyl acetate for 5 minutes and Lead citrate for 2 minutes and viewed with Jeol JEM1400 Plus microscope equipped with Gatan camera.

## Acknowledgments

We wish to thank Yuval Dor and Ayal Ben-Zvi for helpful suggestions and Yael Friedman for EM assistance. This work was supported by the European Research Council (ERC) grant, project VASNICHE (Grant # 322692).

---

**Supplementary Fig. 1.**
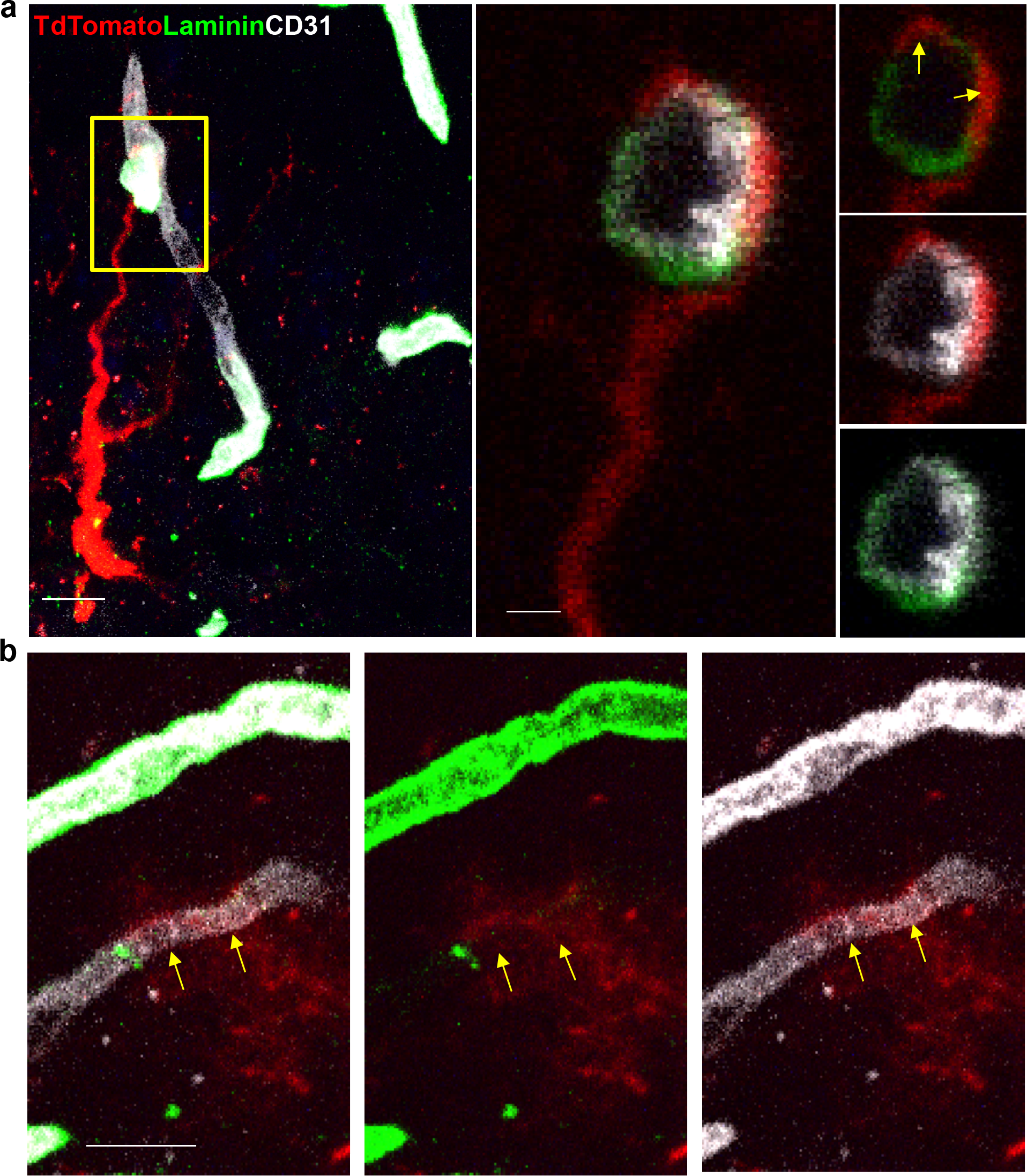
Reduced laminin immunohistochemistry at RGL-BV interface. **a** and **b** Brain sections taken from Gli1-Cre^ER^ mouse crossed to Ai9 reporter, immunostained for CD31 and for the endothelial BM protein Laminin. Scale bar, 10μm. Note reduced laminin staining at the contact points with RGLs (arrows).

